# scGAIN: Single Cell RNA-seq Data Imputation using Generative Adversarial Networks

**DOI:** 10.1101/837302

**Authors:** Mohamed K. Gunady, Jayaram Kancherla, Héctor Corrada Bravo, Soheil Feizi

## Abstract

Single cell RNA sequencing (scRNA-seq) provides a rich view into the heterogeneity underlying a cell population. However single-cell data are usually noisy and very sparse due to the presence of dropout genes. In this work we propose an approach to impute missing gene expressions in single cell data using generative adversarial networks (GANs). By learning an approximate distribution of the data, our approach, scGAIN, can impute dropouts in simulated and real single cell data. The work in this paper discusses how to adopt GAIN training model into the domain of imputing single cell data. Experiments show that scGAIN gives competitive results compared to the state-of-the-art approaches while showing superiority in various aspects in simulation and real data. Imputation by scGAIN successfully recovers the underlying clustering of different subpopulations, provides sharp estimates around true mean expressions and increase the correspondence with matched bulk RNAseq experiments.

## 1 Introduction

Until the recent advent of single-cell methods, RNA sequencing (RNA-seq) was performed in bulk over a cell population to measure its gene expression. Unfortunately, cell-to-cell variation in gene expression within the cell population is lost when RNA sequencing is applied in bulk. Single cell RNA sequencing (scRNA-seq) is designed to capture heterogeneity among different cells within a cell population.

While providing a rich view into the heterogeneity underlying a cell population, data provided by current scRNA-seq technologies are much noisier and sparser than data obtained by bulk RNA-seq. Since the amount of mRNA captured from individual cells is usually low, it will often fail to capture molecules from transcripts with low-to-moderate expression in a considerable number of cells. Genes suffering from such phenomenon are referred to as dropdown genes. Another observed phenomenon is dropout (although recently shown to be less prevalent as originally thought in the literature [8, 17]) where transcripts fail to be captured even in genes with high expression due to technical reasons. Furthermore, because single-cell datasets usually include a heterogenous population of cells, the gene expression distribution in a single gene will show a large number of zeros as some cell types may express the gene and others may not. In other words, a gene expression can be observed as zero in a cell either due to biological (cell types where gene is not expressed) or technical (low sampling or capture failure) reasons. Typical downstream analysis of scRNA-seq data must deal with a highly sparse expression matrix of about 80-90% zeros, arising due to a variety of factors, which can bias the outcome of the analysis if dropouts are not carefully handled. Most clustering, cell type identification and dimensionality reduction methods incorporate methods to handle zeros into their models either through implicit imputation like CIDR [12] or directly modeling the dropout phenomenon like ZIFA [16].

Another tactic is taken by methods that handle zeros to distinguish zeros arising from these factors and impute zeros arising from dropdown and dropout. Many statistical algorithms were developed over the past couple of years that showed effectiveness in recovering dropouts. MAGIC [19], SAVER [10], and scImpute [11] are three leading methods in the field. MAGIC builds Markov transition matrix constructed from cell-to-cell similarity matrix. SAVER builds a Bayesian model to predict the dropouts based on the information of the observed genes. scImpute builds a regression model for each cell and impute only genes with high dropout rates within a cell type by borrowing information from observed genes in that cell type. scImpute requires running cell clustering first to detect cell types before building its imputation model. Although each method adopts a different approach, most of them often demand expensive time and memory resources making scalability a serious drawback of adapting such approaches into more recent large single-cell datasets.

DeepImpute [1] belongs to the category of predictive methods that uses a feed forward neural network to predict a missing gene expression in a cell given the observed values of genes that are highly correlated with that target gene. Recently, a group of methods were proposed to target that problem inspired from the emerging success of generative models. Among these methods are AutoImpute [18], scVI [13] and DCA [4]. scVI uses stochastic optimization and learns cell-specific embeddings using deep neural networks that best explains the observed data, whereas both AutoImpute and DCA uses autoencoders as their interior models. DCA encapsulates data denoising by treating learning the parameters of a zero-inflated negative binomial distribution using an autoencoder network. AutoImpute on the other hand simply trains an autoencoder to learn a latent representation of the data in lower dimension in hope that this representation will focus only on learning the important aspects of the dataset and hence reconstructing a denoised matrix correcting zeros if appropriate. However, both approaches have some limitations. DCA assumes that the dropout distribution follows a predetermined noise distribution which works well for simulated data, it is not clear if these assumptions work well in real data [21]. As for AutoImpute, it works on imputing only the top 1000 differential genes in the dataset, leaving majority of the genes without imputation.

In this paper, we present a novel approach tackling the single-cell data imputation problem using adversarial generative networks (GANs) which we refer to as scGAIN. GANs uses adversarial learning to build a generative model of the data [2, 6, 9]. Generative models are powerful tools that learn the data distribution while having the capability of generating synthesized data points that follows similar characteristics of the existing data. Despite some recent efforts on utilizing the powerful capabilities of GANs into modeling scRNA-seq data [5, 15], they don’t explore the usage of GANs to impute zero values in single-cell data, which causes the underlying models to learn the data distribution including the dropdown and dropout phenomenon so that models generate synthetic that data suffers from the same sparsity. We use capabilities of GANs for accurately and efficiently imputing zero gene expressions due to technical dropdown and dropouts in scRNA-seq data using scGAIN model showed in figure 1c. Experiments show that our model is capable of accurately imputing zeros and providing better estimation of true mean expressions. scGAIN suites large scRNA-seq datasets with thousands and millions of cells which is infeasible for most statistical approaches to deal with. To our knowledge, the is the first work done on using GANs to impute scRNA-seq data which we believe opens the door for new capabilities in imputation.

**Fig. 1.**
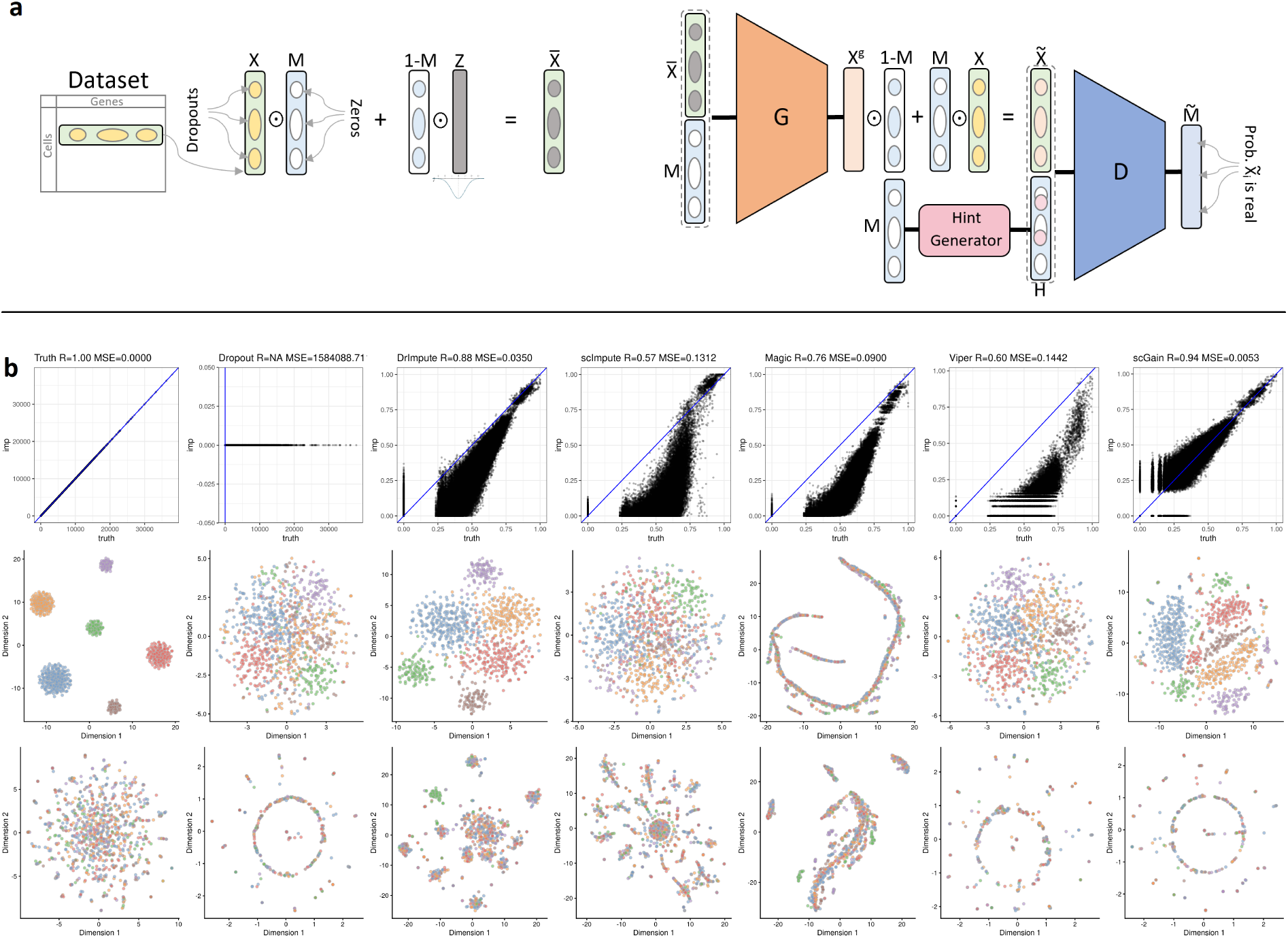
Teaser of scGAIN model and performance for scRNA-seq data imputation. **a)** An illustration of scGAIN’s GAN architecture. **b)** Performance of five different imputation methods (DrImpute, scImpute, Magic, Viper, scGAIN) on simulated dataset (ncells = 20,000, ngenes = 1000) of six groups of cell types with sparsity 85%. Performance is shown in three-fold: 1) Scatter plots showing the correlation and MSE measures between the truth and imputed values. 2) t-SNE visualization of random 1000 cells using the most 100 variable genes from each method. 3) t-SNE visualization of 100 truly non-differential genes.

## 2 Results

### 2.1 scGAIN imputes zeros without adding biases in simulated data

To study the effectiveness of our approach in imputing missing gene expression in a controlled setting where the truth is known, we applied scGAIN to simulated scRNA-seq data generated using Splatter [21]. Splatter is a single-cell data simulator that offers a model simulating dropout in the generated data. It randomly sets some gene expression values to zero following a logistic function based on each gene’s mean expression. We simulated two datasets of three and six cell types. We set Splatter parameters so that the mean sparsity levels of each datasets to be around 65% and 85%, respectively.

Figure 2 shows the results on the first dataset with three cell types. It shows the effect of dropouts in correctly identifying the clusters of the cells. After our approach in scGAIN was used to impute the data, the three clusters were restored and well-separated. Furthermore, it is equally important to examine the effect of imputation on genes that are not differentially expressed across cell types as well as on genes that are differential genes. A proper imputation method should not add any technical bias during the imputation process. We show that by examining the expressions of non-differentially expressed genes before and after imputation. After imputation with scGAIN, the expression levels of those genes remain non-differentially expressed confirming that no bias was introduced into the data by imputation.

**Fig. 2.**
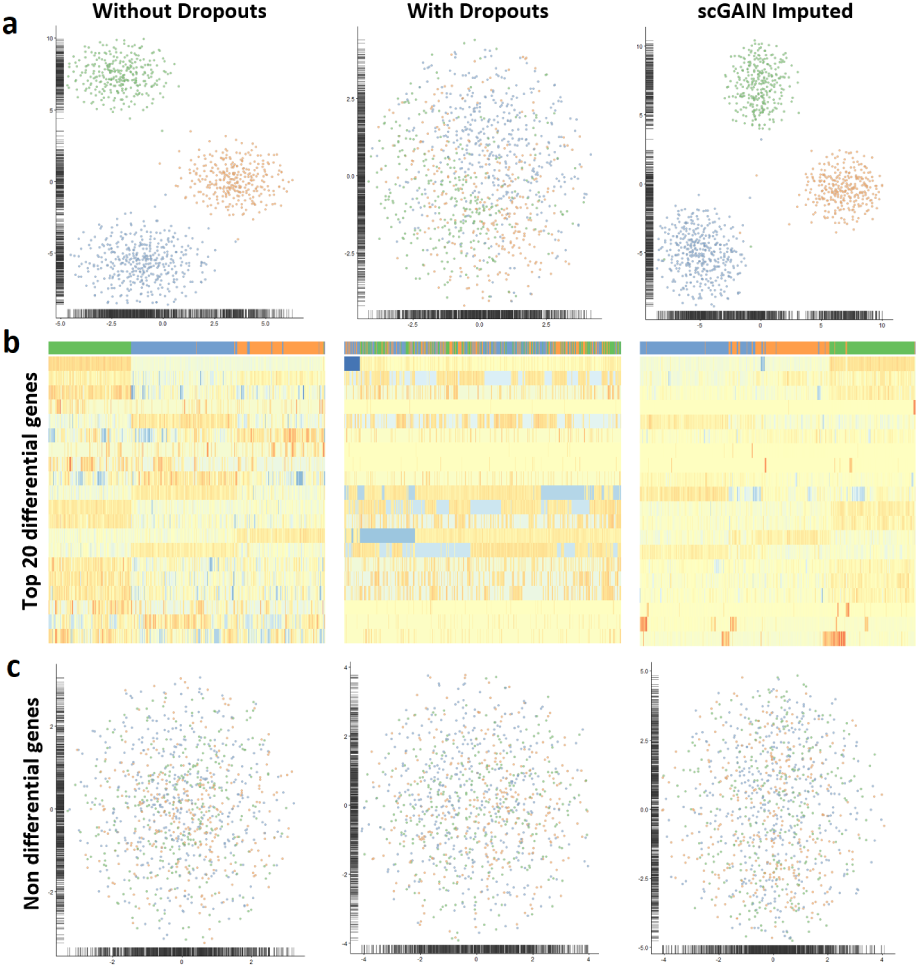
Performance of scGAIN on simulated single-cell dataset of three groups of different cell types. **a**) t-SNE visualization of the three groups of cell types without, with dropouts and after imputation using scGAIN. **b**) Heatmaps of the top 20 differential genes between the cell types. Each column represents a cell with their corresponding cell type color on top. Cells are ordered according to their hierarchical clustering based on the 20 shown genes. **c**) t-SNE visualization of the dataset using the truly non-differential genes only.

Figure 1b shows equivalent results on the second dataset with six cell types and more severe sparsity levels. We show performance of different approaches in three distinct aspects. First, we plot all missing values after imputation versus the true values. We summarize the plot by calculating two metrics: pearson correlation and mean square error (MSE). scGAIN shows best performance in both metrics. Second, we show the effect of imputation in recovering underlying structure of the different cell types, through calculating t-distributed stochastic neighbor embedding (t-SNE) of the true data, raw data without any imputation, and data after imputation by each method. scGAIN successfully recovered the clustering of cells which was relatively distorted in the raw data with the dropouts. Third, t-SNE plots of the same cells using 100 non-differentially expressed genes (across cell types) obtained from the true data. These group of genes remain non-differentially expressed in scGAIN, whereas some methods like DrImpute and Magic showed some undesired clustering.

### 2.2 scGAIN produces stable estimates around the mean with lower variance than other methods

We compare three imputation tools, each representing a category of imputation approaches, on a simulated data (SimData60k). We choose scImpute as a statistical decomposition method, DeepImpute as a prediction method that uses feedforward neural, and our approach (scGAIN) as a generative method. DeepImpute works in a sense as a point estimator of each imputed value. scGAIN on the other hand as a generative model it can give multiple draws of possible values of the missing values following the learnt distribution observed in the population. One way to illustrate that is to see how the individual imputed values are distributed around the true mean expression of the gene in a certain cell type. Figure 3 shows how the individual gene expressions in the imputed matrix (Y-axis) are compared to the corresponding true mean expression (X-axis) used in the simulation. scGAIN shows more stable point estimates around the true mean compared to the other two methods. This high specificity can be desired in imputing data from real experiments since it makes the imputed estimates less susceptible to irrelevant sources of variation and noise. scGAIN consistently gives accurate estimates corresponding to true means even at low expression levels whereas DeepImpute tends to overestimate low and moderate-level expressions.

**Fig. 3.**
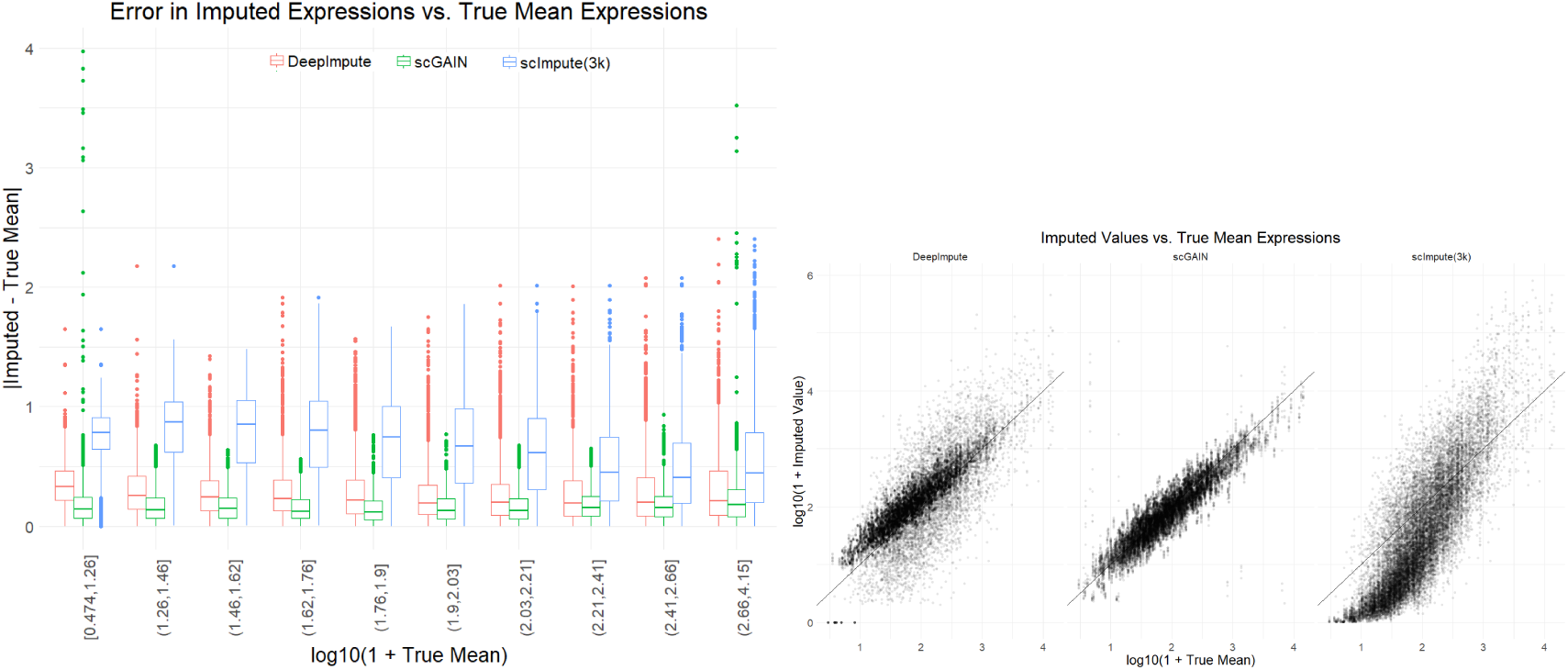
For three imputation methods: scGAIN, scImpute, DeepImpute on simulated data, it shows how the individual gene expressions in the imputed matrix obtained from each method are compared to the corresponding true mean expression used in the simulation. scGAIN shows point estimates around the true mean with smaller variance compared to the other two methods. DeepImpute tends to overestimate low and moderate-level expressions. **(Left)** Box plot of absolute imputation error for each mean expression bin. True mean expressions are divided into bins with equal number of genes. **(Right)** Scatter plot of imputed gene expressions in different cells of one type versus the true mean expression of that gene.

### 2.3 scGAIN increases correspondence with matched bulk expressions of marker genes in PBMC dataset

To evaluate scGAIN on real data and compare its imputation with other methods, we used the PBMC [22] dataset to study the effect of imputation on a set of marker genes for CD4+ vs CD8+ T-cells. First, we determined genes that are differentially expressed between the two cell types from a bulk RNA-seq cell-sorted dataset [7] as the gold standard (by running DESeq [14] and selecting genes with p.value < 0.05). Since not all imputation methods are scalable to impute PBMC in reasonable time, we only include DCA, DeepImpute and MAGIC as a subset for comparison in this analysis. Figure 4a) shows expression of seven marker genes that distinguish between CD4+ and CD8+ cells in bulk, and how their expression levels look like in raw single-cell data and data imputed by scGAIN. ScGAIN helps recovering the correspondance in difference of expressions across the two cell types for each gene.

**Fig. 4.**
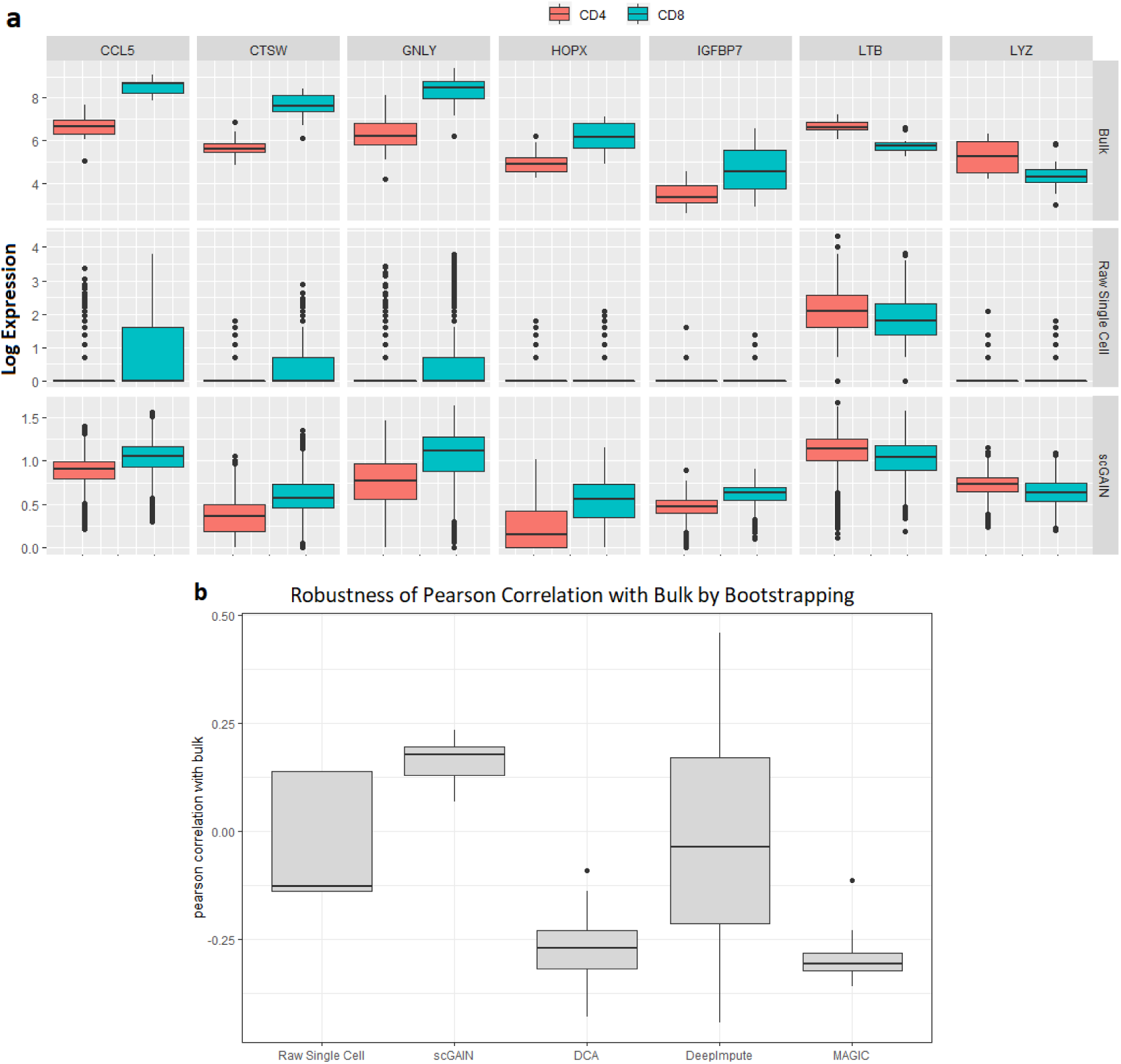
scGAIN increases correspondence with matched bulk expressions of marker genes in PBMC dataset. **a)** shows expression of seven marker genes that distinguish between CD4+ and CD8+ cells in bulk, and how their expression levels look like in raw single-cell data and data imputed by scGAIN. **b)** shows that scGAIN produces the most robustness and the highest correlation among the methods.

Next we wanted to measure the robustness of that correspondence for the top 50 differential genes in bulk with the imputed matrix from each method. Robustness is measured by bootstrapping random 100 cells for 100 iterations, then the Pearson correlation of the difference in log median expressions is calculated for each iteration. Figure 4b) shows that scGAIN produces the most robustness and the highest correlation among the methods.

### 2.4 scGAIN is efficient and scalable to large scRNA-seq experiments

Since scGAIN uses neural network models it scales well to efficiently deal with large datasets which provide enough samples for the network parameters to converge. SimData60k is a typical example of dataset with 60,000 cells. Figure 5 summarizes the running time used by each of the three methods. scGAIN and DeepImpute both have a significant advantage over statistical approaches like scImpute which needs to process the full dataset at once. For instance, scImpute requires as an early step of its pipeline to calculate mutual distances between samples to perform clustering which can be an intensively time and memory consuming process.

**Fig. 5.**
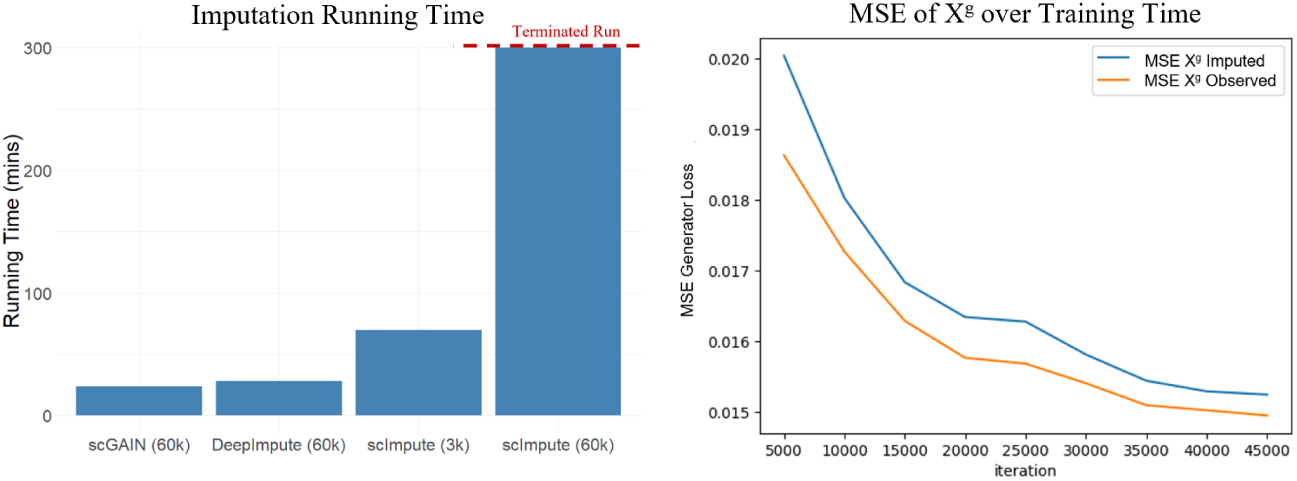
**(Left)** Imputation Running Time for 60,000 cells of Simulated Data (simData60K) used by the three methods. scImpute has two columns: one for imputing a small subset of the dataset of only 3,000 cells, and another for the full dataset. scImpute’s run over the entire 60,000 cells was terminated after running for a full day without finishing. **(Right)** scGAIN’s Generator Loss (MSE) during training iterations. “*X*^*g*^ Imputed” are defined as (1 − *M*) ⊙ *X*^*g*^. “*X*^*g*^ Observed” are defined as *M* ⊙ *X*^*g*^.

Figure 5 shows the convergence of scGAIN over time. Convergence is measured by testing the imputation of 3,000 cells (1,000 from each cell type). MSE is measured between the generated samples and the truth. Two separate curves are shown of the MSE of the imputed portions and the observed portions. The figure shows that both portions of the generated samples are converging, hence the generator is learning the entire sample, rather than just the missing values.

## 3 Methods

### 3.1 Single-Cell Generative adversarial Imputation Nets (scGAIN)

As discussed previously, GANs learns a distribution model for data by learning a network capable of generating samples that resemble the input data. That means the generated samples would carry the same characteristics of the learned data. Hence, if the data itself is sparse, the generator would generate sparse samples as well. Thus, for the purpose of imputation, a rather different model than the standard GAN is required. Another factor to be taken into consideration is that with imputing the missing features in mind as a goal, it is of best interest for the GAN to learn the distribution of the dropdowns and dropouts in the context of the rest of the features. By focusing on learning the distribution of missing features only rather than the entire feature vector, this makes the learning task for GAN more focused and hence speeds up the training convergence. That is why we believe an effective GAN architecture for imputation should have these properties.

Recent efforts have established the use of GANs in imputing missing data in vision and image processing [3, 20]. In this paper, we adopt one of these models from the domain of vision into the biological domain of single-cell RNAseq. As discussed in the introduction section, imputing dropdowns and dropouts in scRNA-seq data can be more challenging. Not only the amount of missing data is significantly larger than it is in the corresponding problem in vision, but also the noise entries are introduced as a function of the missing data. Moreover, the evaluation metric in vision is quite different. When imputing portions of an image, the accuracy of prediction of individual pixels is not that important as long as the final imputed image looks coherent and realistic. While in scRNA-seq single value predictions of individual genes are important. This adds to the challenges when adopting imputation algorithms from vision to our domain of interest. In this paper, we adopt the GAIN model proposed in [20] which was introduced to impute missing portions of pictures. GAIN’s model is to some extent different from a standard GAN. In this section we briefly discuss the main changes that GAIN proposes. Figure 1c shows a diagram of the GAIN model we use for imputing dropouts in single-cell datasets.

#### The Generator

The main goal of the generative model in GAIN is to learn to generate the missing values in a sample *X* = (*x*_1_, *x*_2_, …, *x*_*k*_) for *k* features (genes) rather than focusing on generating the entire feature vector. The generator takes as part of the input a masking vector *M* = (*m*_1_, *m*_2_, …, *m*_*k*_); *m*_*i*_ ∈ {0, 1} dependent on *X* whereas *m*_*i*_ = 0 marks dropdown and dropout genes. The top part in figure 1c shows how the generator input is prepared. The vector 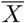 is prepared by filling the missing values in *X* with random noise values from noise vector *Z*. The generator takes a 2k-dimensional vector by concatenating 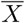 and *M*. The output layer in generator network is a d-dimensional vector *X*^*g*^ where only the imputed portions of it are of interest. By combining the imputed values from *X*^*g*^ with the rest of the observed values *X* we obtain 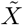 which represent the generated sample from the generator and is fed to the discriminator as input. It should be noted that even though the portions of *X*^*g*^ that doesn’t correspond to dropout genes are replaced by the true observed values in *X* before given to the discriminator as input, the generator still gets penalized by the prediction error in those values as part of the generator’s loss function optimized during training.

#### The Discriminator

Similar to the generator, the discriminator’s model in GAIN is different than standard GAN as well. The main difference is that instead of having a single prediction of whether the entire input sample is real or fake, GAIN’s discriminator gives separate prediction of whether each component 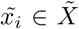 is real or fake given the entire sample. Since the discriminator outputs 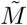 that tends to have zeros for weakly imputed values and ones for true observed values, equivalently it tends to learn the masking vector *M* itself. Yoon et al. showed theoretically in their paper that the discriminator needs as input what they called a *Hint Mechanism*. A hint vector *H* must carry “enough” information about *M* otherwise there would be multiple solutions of G that all can be optimal from the perspective of D. Whilst *H* should not be strictly equal to *M* otherwise the discriminator may trivially converge to outputting *H*. As shown in figure 1c the hint generator component generates a small variation of *M* by setting a few randomly selected values in *M* to 0.5. Components of *M* with 0.5 values say nothing about their corresponding values in 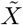 which will be the discriminator job to classify.

#### Objective Function

Along with the changes in the generator and discriminator models from a standard GAN, the loss functions are also slightly different. First, the discriminator no longer takes two sets of samples (real or fake). But rather for a given (imputed) sample, the discriminator should distinguish between observed and imputed portions of that sample. Consequently, the discriminator’s objective function would be for a given *G*:

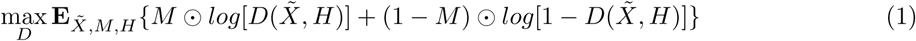

While the generator’s objective function would be for a given *D*:

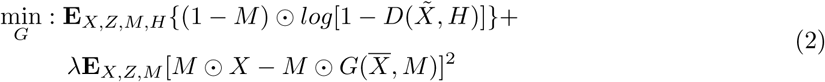

The first term in the loss function represents the cross entropy of the imputed portions of 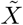. The second regularized term reflects an MSE of the observed portions of 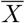.

### 3.2 scGAIN Stratified Training

Throughout our experiments, we noticed that the original GAIN [20] training was not stable when directly applied to single-cell data. In single-cell data is much sparser (samples can have up to 95% zeros) and is not uniformly distributed compared to the standard application of GAIN in image analysis (usually tested against up to 50% sparsity). Figure 6 shows the distribution of zeros in each gene with respect to its mean expression calculated over non-zero observations in the data. Genes with lower mean expressions are more prone to dropdowns.

**Fig. 6.**
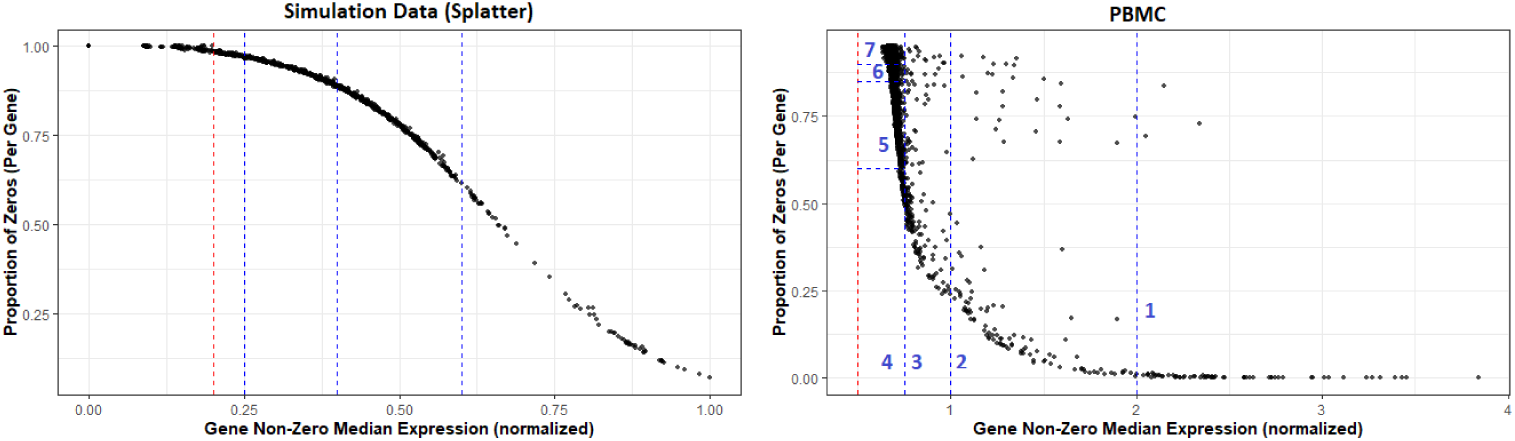
Distribution of zeros in each gene with respect to its mean expression calculated over non-zero observations in simulation data from figure 1 (left) and PBMC (right). Vertical dotted lines represent the different stratification thresholds used during the training on each dataset. Red dotted lines represent the zero cutoff, genes beyond that point were not attempted for imputation. In PBMC, both horizontal and vertical cutoffs are used to stratify the really dense area with low mean expressions, resulting in 7 stratification rounds.

During training, since most zeros in a sample belong to genes with low mean expressions, dropouts in genes of high expressions (which are rarer events) were poorly trained. Those genes barely got any chance to be selected by the hint mechanism to contribute into the discriminator loss function, resulting in a generator with poor accuracy imputing genes with rare dropout events. To remedy that issue, we stratified the training process by capping genes to be masked to a certain threshold. For example, during the first few epochs, the model trains to impute only genes of high mean expression (i.e., dropouts), that controls the number of zeros in the training mask and gives genes with rare sparsity to be learned first. Then gradually the threshold is extended to include more and more genes with higher sparsity rates (i.e., dropdowns). Figure 6 shows vertical lines showing the different stratification thresholds we used during training on simulation dataset from figure 1.

### 3.3 scGAIN Architecture and Parameter Configuration

Both the generator and the discriminator networks have the same structure. Both consist of an input layer of size 2*k* (*k* is the number of genes), then two hidden fully connected layers of 256 then 128 nodes each. Then an output layer of size *k* with a sigmoid activation function to ensure that the output values are in the range [0, 1]. Training is done alternatively between the generator and the discriminator in a fusion of mini-batches of 128 cells at a time. Unless mentioned otherwise, the hint generator would randomly select 10% of the mask vector *H* to mask its values into 0.5. We chose *λ* = 10 as the regularization parameter of MSE term in the loss function of the generator.

### 3.4 Data Normalization

Before data is input to our model, the raw count matrix undergoes several steps of preprocessing. First, the counts are normalized by the total number of UMIs to account for different sequencing depth between cells. Then the normalized data is transformed into log scale. Finally, each cell expression vector is divided by the maximum gene expression present in that cell to ensure that the network takes input values in the range [0, 1].

### 3.5 Imputation Mask Generation

During either training or imputation steps, a mask matrix is required to identify which entries in the input matrix are target for imputation. The mask marks an entry in the data matrix as missing, unless it belongs to a gene of zero or significantly low mean expression determined by a parameter threshold *θ*. Therefore, the mask value of gene *i* in cell *j* is determined as follows:

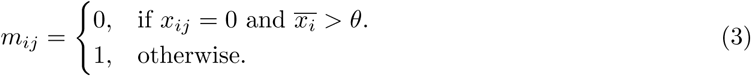

where 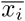 is the mean expression of gene *i* calculated from non-zero values. Note that the threshold *θ* aims at keeping genes of true zeros out of the scope of imputation.

### 3.6 Datasets

#### 1. Simulated Datasets using Splatter

We used Splatter’s R package to generate two simulated datasets of three cell type groups and six cell type groups. In both, we simulated 1000 genes for 20,000 cells. We generated cells such that the proportions the group are 40%, 30% and 30% in the three groups case, and 30%, 20%, 10%, 20%, 10% and 10% in the six groups case. We used parameters dropout.shape = −1, dropout.mid = 3.2 and 5 in R function *splatSimulate()* to add 65% and 85% dropouts into the count matrix, respectively.

#### 2. SimData60k

In this dataset we simulated the gene expression matrix of 2,000 genes for three cell types. We recreated the simulation design and used the same parameters suggested by [11]. log10-transformed read counts were directly generated in the following configuration. For each gene, its mean expression is drawn from a Normal distribution 𝒩 (1.8, 0.5) and its standard deviation is drawn from a Normal distribution 𝒩 (0.6, 0.1). Then 900 genes were randomly selected to be differential across cell types with different 300 genes per each type showing higher expression levels in that cell types than the other two types. We then generate 20,000 cells per cell type using gene means and standard deviations as calculated earlier for each type. Then dropout genes are introduced into both datasets in the same manner. Dropout rates are calculated following a double exponential function of each gene’s mean expression. Finally, zeros are placed into the expression matrix using a Bernoulli distribution based on the calculated dropout rates.

#### 3. PBMC Dataset

This is a dataset of approximately 68,000 cells of peripheral blood mononuclear cells collected from a healthy donor [22]. This dataset is provided publicly on 10x Genomics website.

The data is preprocessed to detect dropouts from genes that consistently show zero expression. This is done by filtering out genes that are zero in 99% of the cells. Then a subset of cells was used as training data in our GAN model. To guarantee that the network is trained on samples with enough data rather than outliers, significantly sparse cells that have less than 2% non-zero genes were excluded from the training data. That leaves a subset of approximately 18,000 cells used as training data to impute about 3,000 genes. It should be noted that this filtering criteria is not mandatory. scGAIN still can accept the full normalized and log-transformed matrix as its input.

## 4 Conclusion

In this paper we propose the application of a generative adversarial network, specifically the GAIN architecture to the problem of imputing the sparse single-cell RNAseq data. This problem manifests some characteristics that are different than the standard GAIN application domain of image analysis. Among those challenges are the severity of the sparsity rates observed in the data (cells can have up to 95% zeros), the distribution of missing values is not uniform over the features, and the absence of ground truth for typical real data. In this work we discussed how we adapted the GAIN model for that purpose through stratified training and mask selection strategies tailored for scRNA-seq data and implemented in scGAIN. We showed that scGAIN was able to correctly impute zeros in simulation data with low and high sparsity without introducing biases into the resulting count matrix. In addition, we showed how scGAIN increases the correspondence of differential expression analysis between PBMC and bulk RNAseq for CD4+ and CD8+ T-cell marker genes and increases the robustness of differential expression statistic estimates obtained from scRNA-seq data.

## A Generative Adverserial Networks

A standard GAN [6] consists of two network components: Generator and Discriminator. The Generator network attempts to map samples from a low-dimensional high entropy distribution such as a Normal distribution, into a low-dimensional manifold within the high-dimensional space from which data is observed. The Discriminator network attempts to differentiate between samples from the real dataset and fake samples generated by the generator. Both components are trained jointly in an adversarial fashion where a min-max optimization problem forms a game between the generator and the discriminator. The generator’s goal is to generate fake samples that are very close to the real samples such that the discriminator unable to distinguish between them, while the discriminator’s goal is to learn the real data well that it would be able to accurately distinguish fake from real samples. When training converges, the generator would have reached a point that it can generate fake samples resembling the real distribution such that the well-trained discriminator is no longer able to distinguish between a sample present in the dataset and a sample generated by the generator.

Figure 7 shows a diagram of a standard GAN. Generator G takes low-dimensional random noise vector **Z** = (*z, z, …, z*_*l*_); *z*_*i*_ ∼ 𝒩 (0, 1) as input and generates a sample 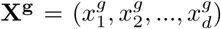. Discriminator D takes either a real sample **X** or a fake sample *X*^*g*^ and outputs whether the input sample is real of fake. A well-trained generator can generate random samples that follows the same distribution of **X** by drawing random values of **Z**. The training of a GAN typically tries to solve the optimization problem

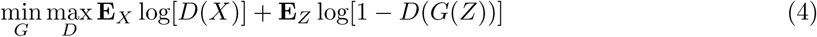

**Fig. 7.**
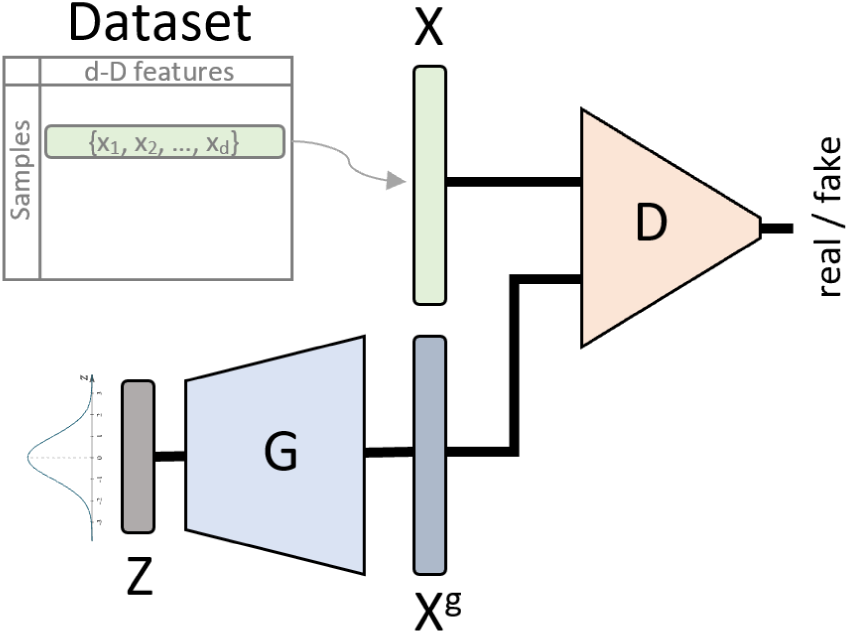
Diagram of a standard GAN architecture. Generator G takes low-dimensional random noise vector Z as input and generates a d-D sample *X*^*g*^. Discriminator D takes either a real sample X or a fake sample *X*^*g*^ and outputs whether the input sample is real of fake.

## B Homogenous Cell Population from PBMC are Zero-Inflated

Recent work argues that droplet scRNA-seq data, for example the PBMC data, is not zero-inflated when considering technical replicates [17]. This would imply observed zeros are the result of dropdown and cell-type heterogeneity. In order to confirm that the observed zero-inflation in single-cell raw data is not strictly the result of having heterogenous population of cells, we subset one subpopulation (CD19+ B) from PBMC. Figure 8 shows that zero-inflation fits data better even when looking at a homogenous cell population of CD19+ B cells.

**Fig. 8.**
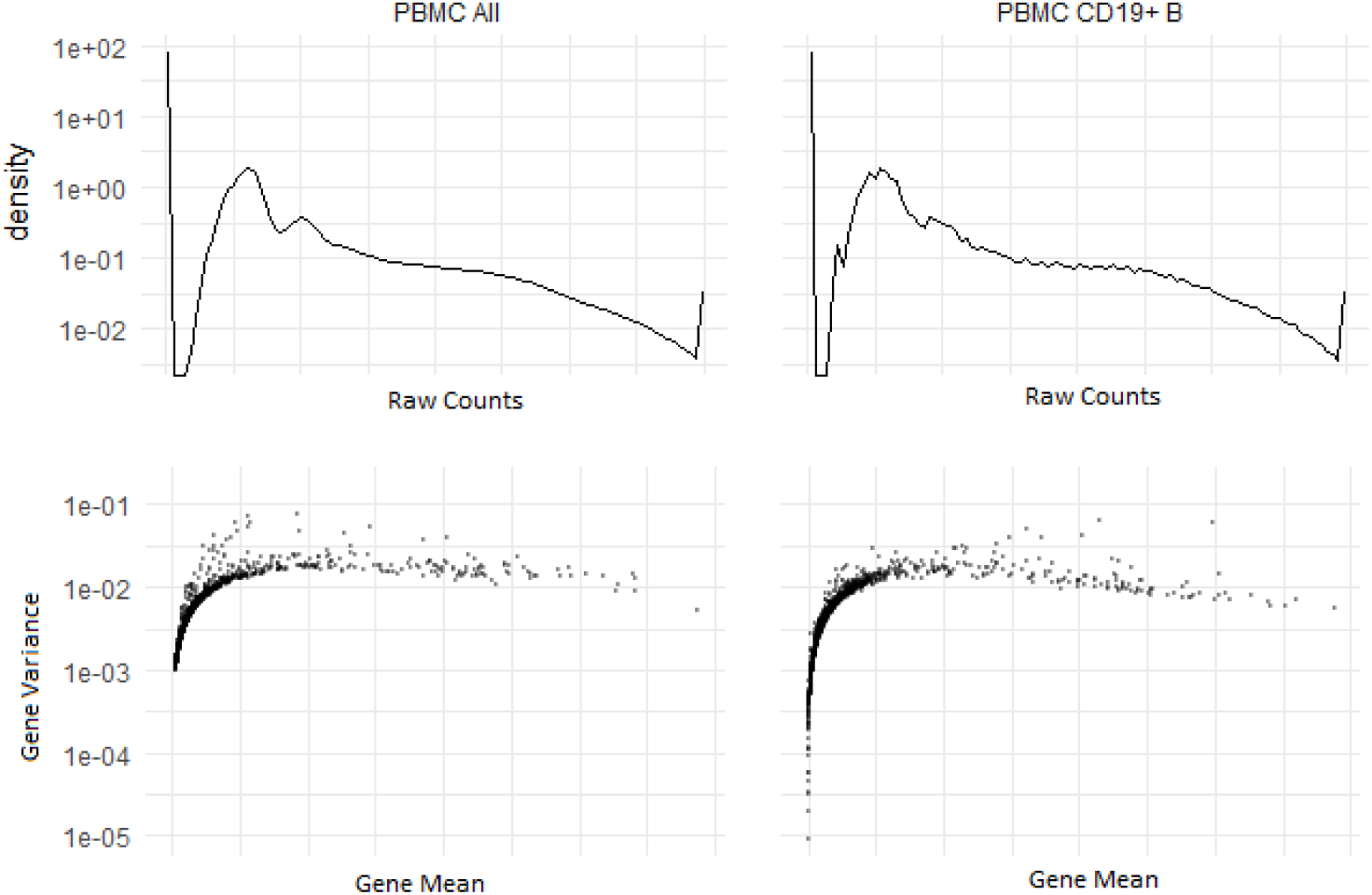
Plot showing the distribution of raw counts and mean-variance expressions of genes from PBMC dataset. It shows that the data is zero-inflated with heterogenous cell population (PBMC All) as well as homogenous cell population (PBMC CD19+ B).

